# Bright photoactivatable fluorophores for single-molecule imaging

**DOI:** 10.1101/066779

**Authors:** Jonathan B. Grimm, Brian P. English, Anand K. Muthusamy, Brian P. Mehl, Peng Dong, Timothy A. Brown, Zhe Liu, Timothée Lionnet, Luke D. Lavis

## Abstract

Small molecule fluorophores are important tools for advanced imaging experiments. The development of self-labeling protein tags such as the HaloTag and SNAP-tag has expanded the utility of chemical dyes in live-cell microscopy. We recently described a general method for improving the brightness and photostability of small, cell-permeable fluorophores, resulting in the azetidine-containing “Janelia Fluor” (JF) dyes. Here, we refine and extend the utility of the JF dyes by synthesizing photoactivatable derivatives that are compatible with established live-cell labeling strategies. These compounds retain the superior brightness of the JF dyes but their facile photoactivation enables improved single-particle tracking and localization microscopy experiments.

---

Small-molecule fluorophores are brighter than fluorescent proteins and remain a crucial element of modern microscopy methods.^1,2^ The development of new protein-specific labeling strategies, such as the self-labeling tag concept pioneered by Johnsson,^3,4^ enables the formation of fluorescent bioconjugates inside living cells: bright, membrane-permeable synthetic dyes passively diffuse into cells where they form covalent bonds with their genetically-encoded cognate protein fused to a factor of interest. Self-labeling tags thus combine simple genetic encoding—one of the main advantages of fluorescent proteins—with the favorable photophysics of organic fluorophores. Building upon these sophisticated attachment techniques, we recently reported replacement of ubiquitous dimethylamino groups in classic fluorophores with four-membered azetidine rings as a general strategy for improving the brightness and photostability of small, cell-permeable fluorophores.^5^ These “Janelia Fluor” (JF) dyes are excellent labels for live-cell imaging, especially in single-molecule tracking experiments where the improved brightness and photon counts allowed longer observations and better localization of individual fluorescent conjugates. We now report photoactivatable (PA) versions of JF_549_ and JF646, demonstrate their compatibility with existing live-cell labeling strategies, and show their utility in single-molecule tracking and super-resolution imaging.

JF_549_ and JF_646_ are fully N-alkylated rhodamine dyes and cannot be caged using N-acylation with standard photolabile groups as can other rhodamine dyes.^6^ Instead, we utilized a caging strategy serendipitously discovered by Hell and coworkers, in which treatment of rhodamine dyes with oxalyl chloride and diazomethane generates a spirocyclic diazoketone that is colorless and nonfluorescent.^7,8^ Activation of these diazoketone dyes with short-wavelength light yields a fluorescent species as the major product. Although diazoketone-caged dyes have been employed as antibody labels, this type of photoactivatable dye has not been incorporated into self-labeling tag systems nor used in live-cell single-particle tracking experiments.

To test the compatibility of this caging strategy with the azetidinyl Janelia Fluor dyes, we first prepared the photoactivatable JF_549_ (PA-JF_549_, **2**) in good yield from JF_549_ (**1**^5^, **Fig. 1a**). We then tested the photoactivation of this molecule in the presence of water, whereas previous experiments had only examined this photochemical reaction in methanol.^7,8^ Surprisingly, the major product formed was not the expected phenylacetic acid dye 3 but the methyl-substituted JF_549_ (**4**, **Fig. 1a**). This result is likely due to facile photoinduced decarboxylation^9^ of the initial photochemical product **3** (**Supplementary Note**). Nevertheless, the resulting dye **4** is a highly fluorescent molecule with similar spectral properties to the parent JF_549_ (**1**; **Fig. 1b**). As reported before,^5^ fluorophore **1** exhibits an absorption maximum (λ_max_) of 549 nm, extinction coefficient (ε) of 1.01 × 10^5^ M^−1^cm^−1^, emission maximum (λ_em_) of 571 nm, and a quantum yield (ϕ) of 0.88. Dye **4** gave λ_max_/λ_em_ = 551 nm/570 nm and retained 88% of the brightness of **1** (ε = 9.83 × 10^4^ M^−1^cm^−1^; ϕ = 0.80).

**Figure 1.**
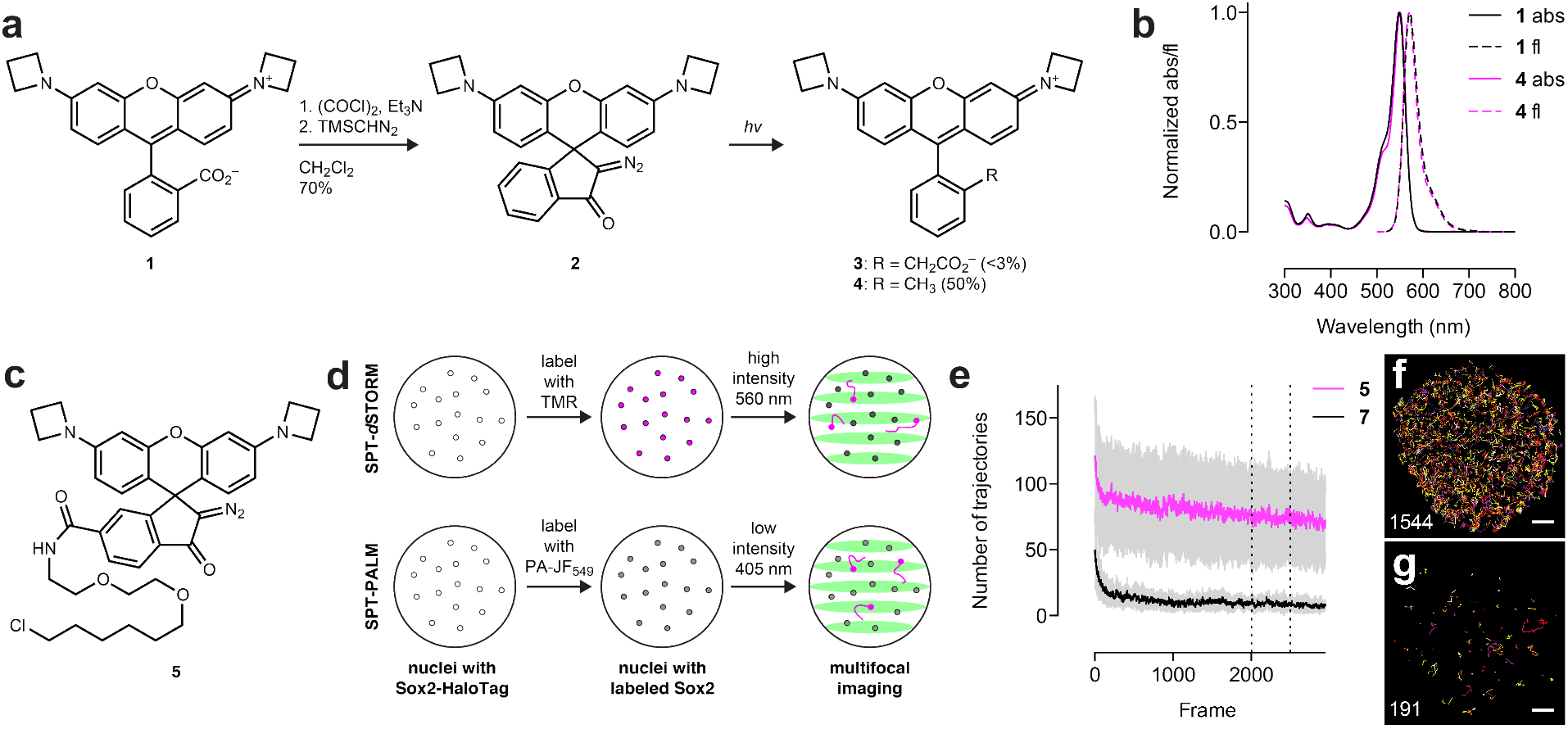
Synthesis, characterization, and utility of photoactivatable Janelia Fluor 549 (PA-JF_549_). (**a**) Synthesis of PA-JF_549_ (**2**) from JF_549_ (**1**). Photoconversion (365 nm) in the presence of water yields methyl-substituted JF_549_ (**4**) as the major product with only trace amounts of expected product **3**. (**b**) Normalized absorption (abs) and fluorescence emission (fl) of **1** and **4**. (**c**) Structure of PA-JF_549_-HaloTag ligand (**5**). (**d**) Cartoon showing experimental workflow of SPT-*d*STORM experiment (top) and SPT-PALM experiment (bottom). ES cells expressing Halo Tag–Sox2 were labeled to saturation with PA-JF_549_ ligand **5** or TMR-HaloTag ligand **7** and imaged on a multifocus microscope, which allows simultaneous imaging of 9 focal planes across an axial depth of ~4 μm. (**e**) Plot of the number of single molecule trajectories measured per frame in ES cell expressing HaloTag–Sox2 labeled with PA-JF_549_ ligand **5** (SPT-PALM mode, magenta) or the commercial TMR-HaloTag ligand (**7**; SPT-*d*STORM mode, black); n = 5 cells for each ligand; s.d. shown in gray. (**f–g**) Image of cumulative single-particle tracks imaged for frames 2000–2500 (between dashed lines in e; only trajectories observed in >5 successive frames shown); lower left: number of trajectories measured; lower right: scale bars: 2 μm. (**f**) Cell labeled with PA-JF_549_ ligand **7** (1544 trajectories). Cell labeled with standard TMR ligand 5 (191 trajectories).

Given the brightness of the resulting photoactivated fluorophore we then synthesized the HaloTag ligand^10^ of PA-JF_549_ (**5**, **Fig. 1c, Supplementary Note**). Labeling of HaloTag protein with **5** either *in vitro* or in live cells gave conjugates with low background absorption and fluorescence that could be activated by one- or two-photon illumination (**Supplementary Fig. 1a**, **Supplementary Videos 1–2**). We found attachment to the HaloTag improves the yield of the desired fluorescent product compared to the free dye (**Supplementary Note**) and the absorptivity of the photoactivated conjugate was similar in magnitude to HaloTag protein labeled with the standard JF_549_-HaloTag ligand **6** (**Supplementary Fig. 1b,c**). We directly compared the performance of the PA-JF_549_-HaloTag ligand (**5**) to the commercially available tetramethylrhodamine (TMR) HaloTag ligand (**7**, **Supplementary Fig. 1b**)^10^ in 3D single-particle tracking (SPT) experiments in mouse embryonic stem (ES) cells expressing HaloTag-Sox2 fusions using a multifocus microscope (MFM) setup (**Fig. 1d**).^11^ To achieve the sparse labeling required for SPT with the commercial TMR ligand 7, we used high initial illumination power to shelve molecules in a dark state based on live-cell *direct* stochastic optical reconstruction microscopy (dSTORM).^12^ Although this SPT-dSTORM technique was successful, it gave a relatively low number of single-molecule tracks (=10446; **Fig. 1e,f**, **Supplementary Fig. 1d**, **Supplementary Video 3**) due to the difficulty in controlling the on-state of the fluorophore and the irreparable loss of dyes under the high initial excitation power.^13^ The use of a photoactivatable fluorophore such as PA-JF_549_ ligand **5** overcomes this limitation, allowing controlled activation with low amounts of 405 nm light from the entire initial pool of PA-JF_549_. This SPT based on photoactivated localization microscopy (SPT-PALM) results in many more single-molecule tracks per cell (=40324; **Fig. 1e,g**, **Supplementary Video 3**). Moreover, the PA-JF_549_ ligand retained the improved brightness of the standard Janelia Fluor 549 dye resulting in higher localization precision. The commercial TMR ligand showed a mean photon yield per particle per frame of 103 yielding a mean localization error (σ) of 54 nm whereas the PA-JF_549_ ligand (**5**) gave more photons/particle/frame (mean = 162, σ = 45 nm; **Supplementary Fig. 1e,f**).

We then attempted two-color single-particle tracking, an experiment that has been stymied by the requirement of sparse labeling of two separate molecular species. Control of two fluorophores under the dSTORM imaging mode is difficult, requiring matching of the blinking phenomena with two separate dyes and excitation powers. Additionally, the labeling kinetics of the HaloTag and SNAP-tag differ by 1000-fold^10,14^ and the respective ligands exhibit dissimilar membrane permeability, making it challenging to control the labeling of two populations of proteins at levels low enough to image single molecules. We reasoned that the use of the same diazoketone caging strategy on two spectrally distinct dyes could allow sparse photoactivation of both labels with similar efficiency, thus facilitating two-color experiments. Accordingly, we synthesized the HaloTag and SNAP-tag ligands based on a photoactivatable derivative of the farred Janelia Fluor 646 (**8** and **9**, PA-JF646, **Fig. 2a**, **Supplementary Note**). Like the PA-JF_549_-HaloTag ligand (**5**), the PA-JF_646_-HaloTag ligand (**8**) showed low background fluorescence and facile activation with blue light (**Supplementary Video 4**) and conjugation of compound 8 to the HaloTag protein increased the yield of the desired fluorescent photoproduct (**Supplementary Note**).

**Figure 2.**
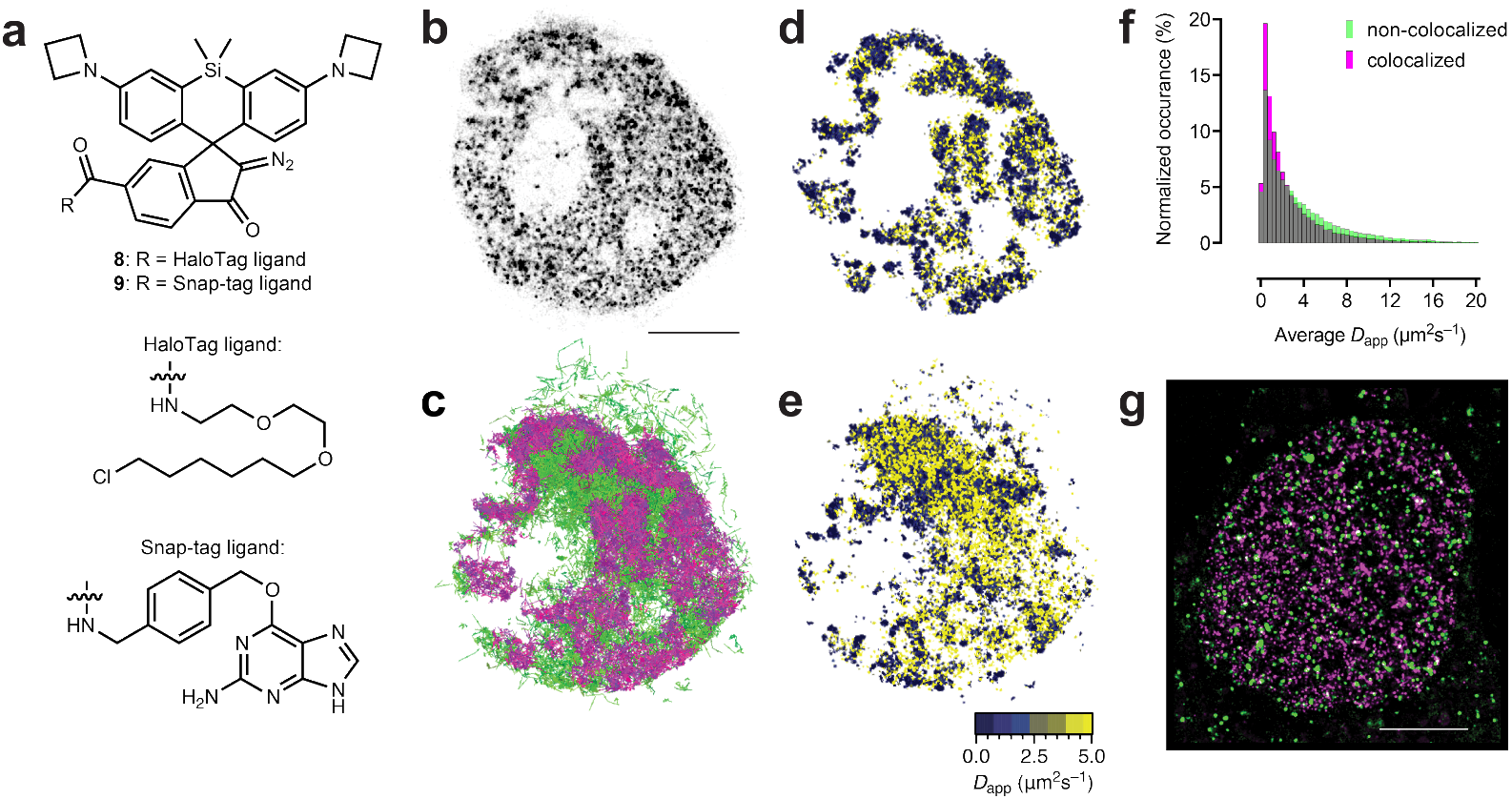
Multicolor imaging using photoactivatable Janelia Fluor 646 (PA-JF_646_). (**a**) Structures of PA-JF_646_-HaioTag ligand (**8**) and PA-JF_646_-SNAP-tag ligand (**9**). (**b–f**) Simultaneous two-color SPT-PALM experiment in a live ES cell expressing histone H2B–SNAP-tag labeled with **9** and Sox2-HaloTag labeled with PA-JF_549_-HaloTag ligand (**5**). (**b**) PALM image histone H2B–SNAP-tag; scale bar: 5 μm. (**c**) Singleparticle trajectories of Sox2–HaloTag that are colocalized with histone H2B–SNAP-tag (6272 trajectories, magenta) or non-colocalized with histone H2B–SNAP-tag (7081 trajectories, green). (**d**) Apparent diffusion coefficient map of colocalized fraction of Sox2. (**e**) Apparent diffusion coefficient map of non-colocalized fraction of Sox2. (**f**) Histogram of normalized occurrence vs. apparent diffusion coefficient calculated for each step in the colocalized and non-colocalized Sox2 trajectories. (**g**) Overlay of the PALM image of Htt84Q–mEos3.2 clusters with the PALM image of histone H2B–HaloTag labeled with 8. The PALM images were simultaneously recorded and are each composed of 10,000 consecutive frames. The 76,341 detected mEos3.2 molecules (green) and the 190,710 PA-JF_646_ molecules (magenta) are displayed according to their localization full-with at half-maximum. The median number of detected photons per mEos3.2 molecule was 114.3, and the median number of detected photons per PA-JF_646_ molecule was 805.3. The median localization error for mEos3.2 was 35.8 nm, and for PA-JF_646_ was 20.6 nm. Scale-bar: 5 μm.

In previous work^5^ we demonstrated two-color single-molecule imaging using the Janelia Fluor dyes by first performing *d*STORM^12^ on histone H2B labeled with JF_549_ followed by tracking of a diffusive tracer TetR labeled with JF_646_. Although this experiment demonstrated the brightness of the JF dyes and the feasibility of two-color experiments, it relied on the fact that histone H2B was largely static until the subsequent single-particle tracking. A more rigorous experiment would be simultaneously tracking of both markers, generalizing our live-cell single-molecule tracking scheme to any pair of proteins. As above, the transcription factor Sox2 was expressed as a fusion with HaloTag protein and labeled with PA-JF_549_ ligand 5. We coexpressed histone H2B as a fusion with the SNAP-tag and labeled this population with PA-JF_646_-SNAP-tag ligand (**9**); these photoactivatable dyes allowed simultaneous tracking of both H2B and Sox2. We generated a map of histone H2B location using a standard PALM analysis (**Fig. 2b**) and used it to define Sox2 trajectories that were either colocalized or not colocalized with this chromatin marker (**Fig. 2c**). As expected, the molecules of Sox2 that were colocalized with histone H2B exhibited slower diffusion coefficients than the non-colocalized fraction (**Fig. 2d–f**, **Supplementary Fig. 2**).

Finally, we investigated the PA-JF dyes as labels for photoactivatable localization microscopy (PALM). Although a few self-labeling tag ligands have been used in localization microscopy,^8,15,16^ previously reported molecules exhibit relatively short emission maxima, and are thus incompatible with established PALM labels such as photoconvertible fluorescent proteins. Based on previous work with a caged Si-rhodamine,^6^ we reasoned that PA-JF_646_ would be red-shifted enough to be useful for two-color photoactivated localization microscopy (PALM)^17^ with the genetically encoded photoconvertable protein mEos3.2.^18^ We expressed the mutant Huntingtin protein Htt-94Q as a fusion protein with mEos3.2 and histone H2B as a fusion with the HaloTag, labeling with PA-JF_646_-HaloTag ligand **8**. After labeling, fixation, and two-color PALM (**Fig. 2g**), we observed that histone H2B (magenta) and Htt-94Q aggregates (green) only rarely overlap (i.e., few white spots), which supports the hypothesis that the aggregates formed by expanded polyglutamine domains displace chromatin structures in the nucleus. Notably, the JF646 dye was significantly brighter than mEos3.2, emitting seven-fold more photons (median detected photons = 805) relative to the fluorescent protein (median detected photons = 114).

In conclusion, we report photoactivatable versions of the bright, photostable Janelia Fluor dyes. These fluorophores retain the superior properties of the JF dyes but have the added benefit of photoactivation, allowing sophisticated single-particle tracking experiments and facile superresolution microscopy. For single-particle tracking experiments they circumvent the problems associated with sparse activation of standard fluorophores and differential labeling efficiencies. For super-resolution microscopy, they allow facile labeling and activation of the latent fluorophores. In particular, JF_646_ is the first far-red photoactivatable fluorophore compatible with live-cell labeling using the HaloTag or SNAP-tag systems, allowing multicolor single-particle tracking experiments and super-resolution microscopy with photoconvertible fluorescent proteins. Unlike traditional photoactivatable xanthene dyes that contain large, poorly soluble caging groups,^6^ we expect these small and bright photoactivatable labels to be compatible with many different labeling strategies, therefore extending the boundaries of single-molecule imaging in live and fixed cells. Beyond single molecule imaging, these versatile labels are simple to use and should provide a favorable alternative to photoconvertible fluorescent proteins in any imaging experiment where photoactivation is used to highlight a specific cell or cellular region.

## Methods

### Chemical Synthesis

Experimental details and characterization for all novel compounds and subsequent spectroscopy and photochemistry experiments can be found in the Supplementary Note.

### UV–Vis and Fluorescence Spectroscopy

Spectroscopy was performed using 1-cm path length quartz cuvettes. All measurements were taken at ambient temperature (22 ± 2 °C). Absorption spectra were recorded on a Cary Model 100 spectrometer (Varian). Fluorescence spectra were recorded on a Cary Eclipse fluorometer (Varian). Absolute fluorescence quantum yields (ϕ) for all fluorophores were measured using a Quantaurus-QY spectrometer (model C11374, Hamamatsu).

### Cell Culture

Mouse D3 ES cells (ATCC) were maintained on 0.1% w/v gelatin coated plates in the absence of feeder cells. The ES cell medium was prepared by supplementing knockout Dulbecco’s modified eagles media (DMEM, Invitrogen) with 15% v/v fetal bovine serum (FBS), 1 mM glutamax, 0.1 mM nonessential amino acids, 1 mM sodium pyruvate, 0.1 mM 2-mercaptoethanol, and 1000 units of leukemia inhibitory factor (LIF; Millipore). Cells were regularly tested for mycoplasma contamination by the Janelia Cell Culture Facility.

### Plasmid Construction

Sox2 and histone H2B cDNA were amplified from ES cell cDNA libraries. Htt-94Q cDNA was obtained from Addgene (Plasmid #23966). The full-length cDNAs were cloned into the Piggybac transposon vector (PB533A-2, System Biosciences) or a modified Piggybac transposon vector with PuroR. The sequence for HaloTag (Promega) or mEOS3.2 (Addgene: Plasmid #54525) was ligated in-frame with the cDNA of the desired proteins at the N-terminus (HaloTag–Sox2) or C-terminus (histone H2B–HaloTag, histone H2B–SNAP-tag, and Htt-94Q–mEOS3.2-NLS).

### Stable Cell Line Generation

Stable cell lines were generated by co-transfection of Piggybac transposon vector with a helper plasmid that over-expresses Piggybac transposase (Super Piggybac Transposase, System Biosciences). At 48 h post-transfection, ES cells were subjected to neomycin or puromycin (Invitrogen) selection. Transfection was conducted by using the Nucleofector Kit for Mouse Embryonic Stem Cells (Lonza).

### Cell Labeling Strategy and Preparation for Imaging

One day before imaging, ES cells were plated onto a cover slip pre-coated with IMatrix-511 (Clontech). Imaging was performed in the ES cell imaging medium, which was prepared by supplementing FluoroBrite medium (Invitrogen) with 10% v/v FBS, 1 mM glutamax, 0.1 mM nonessential amino acids, 1 mM sodium pyruvate, 10 mM HEPES (pH 7.2–7.5), 0.1 mM 2-mercaptoethanol, and 1000 units of LIF (Millipore). For PA-JF_646_ or PA-JF_549_ labeling, cells were incubated with PA-JF_549_-HaloTag ligand (**5**) or PA-JF_646_-HaloTag ligand (**9**) at a final concentration of 100 nM for 1 h. For the 2-color SPT-PALM live-cell tracking experiments, labeled cells were washed with ES cell imaging medium (3×) before imaging. For the 2-color fixed-cell PALM imaging experiments, labeled cells were washed with PBS (4×), fixed in 4% w/v paraformaldehyde for 10 min and washed with PBS (3×). The final PALM imaging was performed in PBS solution.

### Two-color SPT-PALM live-cell tracking experiments

ES cells expressing both HaloTag–Sox2 fusions labeled with PA-JF_549_-HaloTag ligand (**5**) and SNAP-tag–histone H2B fusions labeled with PA-JF_646_-SNAP-tag ligand (**9**) were tracked simultaneously using a custom-built 3-camera microscope.^19^ Two iXon Ultra EMCCD cameras (DU-897-CS0-BV and DU-897U-CS0-EXF, both cooled to −80 °C, 17MHz EM amplifiers, pre-amp setting 3, Gain 400) were synchronized using a National Instruments DAQ board (NI-DAQ-USB-6363) at a frame time of 10 ms. 5 ms stroboscopic excitations of a 555 nm laser (CL555-1000-O with TTL modulation, CrystaLaser) and a 639 nm laser (Stradus 637–140, Vortran) were synchronized to the frame times of the two respective cameras via LabVIEW 2012 (National Instruments). The two lasers stroboscopically illuminated the sample using peak power densities of around 20 kW/cm^2^ using HiLo illumination of the nucleus The PA-JF_549_ and PA-JF_646_ labels were photoconverted by 100 μs long excitation pulses of 407 nm light (50 W/cm^2^) every second. During the course of image acquisition, the pulse length was increased to 200 μs long pulses. During imaging, cells were maintained at 37 °C and 5 % CO_2_ using a Tokai-hit stage top incubator and objective heater.^19^

### Two-color fixed-cell PALM imaging acquisition

ES cells expressing both Htt94Q–mEOS3.2 and HaloTag–histone H2B labeled with PA-JF646-HaloTag ligand (8) were imaged using the previously described custom-built 3-camera microscope at a frame time of 50 ms and a constant illumination power density of around 20 kW/cm^2^ for both 555 nm and 639 nm excitation lasers. mEOS3.2 and PA-JF_646_ were photoconverted by 100 μs long excitation pulses of 407 nm light (100 W/cm^2^) every second.

### 3D SPT-*d*STORM and SPT-PALM tracking experiments

Fluorescently tagged HaloTag–Sox2 molecules labeled either with PA-JF_549_-HaloTag ligand (**5**) or with TMR-HaloTag ligand (**7**) were tracked in live ES cells in 3D using a custom-built multifocus microscope.^11^ The fluorescence from nine focal planes was simultaneously recorded using an iXon Ultra EMCCD camera (DU-897U-CS0-#BV, 17MHz EM amplifiers, pre-amp setting 1, Gain 300) at a frame time of 30 ms.

### 2-color super-resolution imaging and SPT-PALM tracking image analysis

For simultaneous 2-camera imaging and tracking, the two 16-bit TIFF stacks were registered using the similarity (2d) transformation model using a descriptor-based Fiji plugin.^20^ Super-resolution images were rendered using the software package Localizer by Dedecker *et al*.^21^ with 8-way adjacency particle detection, 20 GLRT sensitivity, and a PSF of 1.3 pixels. The following settings were chosen for particle track linking: 5 pixel maximum jump distance, 3-frame minimum track length, and 15 GLRT sensitivity. Resulting tracks were then exported as text files, and diffusion mapping was performed with code written in Igor Pro 6.36 (WaveMetrics). The code calculates local apparent diffusion coefficients evaluated in 20 nm by 20 nm grids from the mean square displacements over the frame-time timescale.^22^

### Multifocus image processing

We assembled 3D stacks by aligning the nine simultaneously obtained focal planes on top of one another using bead calibration data as described previously.^11^ For 3D particle tracking we imported the 16-bit TIFF stack into DiaTrack 3.04 Pro,^23^ which identifies and fits the intensity spots with 3D Gaussian function matched to a pre-determined PSF. The following settings were chosen for 3D particle tracking: Subtract background, Filter data of 1.05, PSF of 1.3 pixels, remove dim of 15, and remove blurred of 0.05. Resulting 3D tracks were exported with code written in Igor Pro 6.36 as one text file containing frame numbers, as well as x, y, and z-coordinates of all detected points. We plotted a map of all detected particle locations in the x-y plane, color-coded for height (z), and calculated histograms of detected number of particles over the course of 3D SPT-PALM data acquisitions. Local diffusion mapping in the x-z plane was performed with code that calculates local apparent diffusion coefficients evaluated in 20 nm by 20 nm grids and displays the diffusion map as an x–z projection. Integrated fluorescence intensities from particles detected in the central two focal planes (multifocal plane 4 and 5) were calculated and converted to photon counts using analysis routines written in Igor Pro version 6.36.^5^ Localization errors were calculated using equation (6) in Mortensen *et al*.^24^

## Acknowledgement

We thank Wesley Legant and Eric Betzig (Janelia) for contributive discussions and Adam Berro and Eric Schreiter (Janelia) for the purified HaloTag protein.

---

**Supplementary Figure 1.**
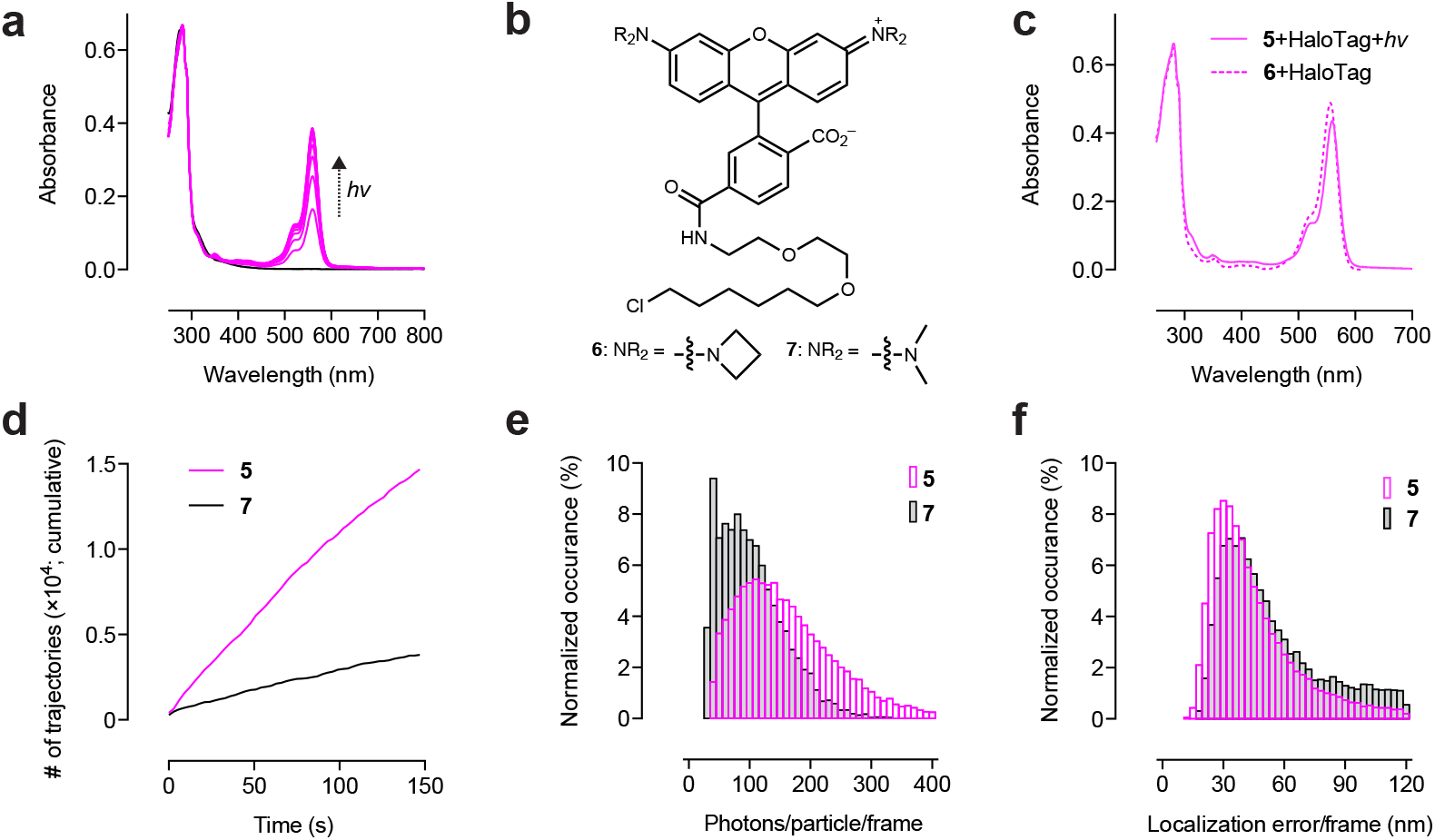
Performance of PA-JF_549_-HaloTag ligand. (**a**) Absorbance spectrum of PA-JF_549_-HaloTag ligand (**5**) bound to HaloTag protein before (black line) and after photoactivation (365 nm; magenta lines). (**b**) Structure of JF_549_-HaloTag ligand (**6**) and TMR-HaloTag ligand (**7**). (**c**) Comparison of the absorbance spectra of PA-JF_549_-HaloTag ligand (**5**) bound to HaloTag protein and then exhaustively photoactivated (solid line) with the absorbance spectrum of JF_549_-HaloTag ligand (**5**) bound to HaloTag protein (dashed line). (**d–f**) Statistics from 3D tracking experiment in ES cells expressing HaloTag–Sox2 and labeled with **5** or **7** (**Fig. 1d–g**). (**d**) Plot of the cumulative number of trajectories vs. time using labels **5** (magenta) or **7** (black). (**e**) Histogram of normalized occurrence vs. photons/particle per frame using labels **5** (magenta) or 7 (black). (**f**) Histogram of normalized occurrence vs. particle localization per frame using labels **5** (magenta) or **7** (black).

**Supplementary Figure 2.**
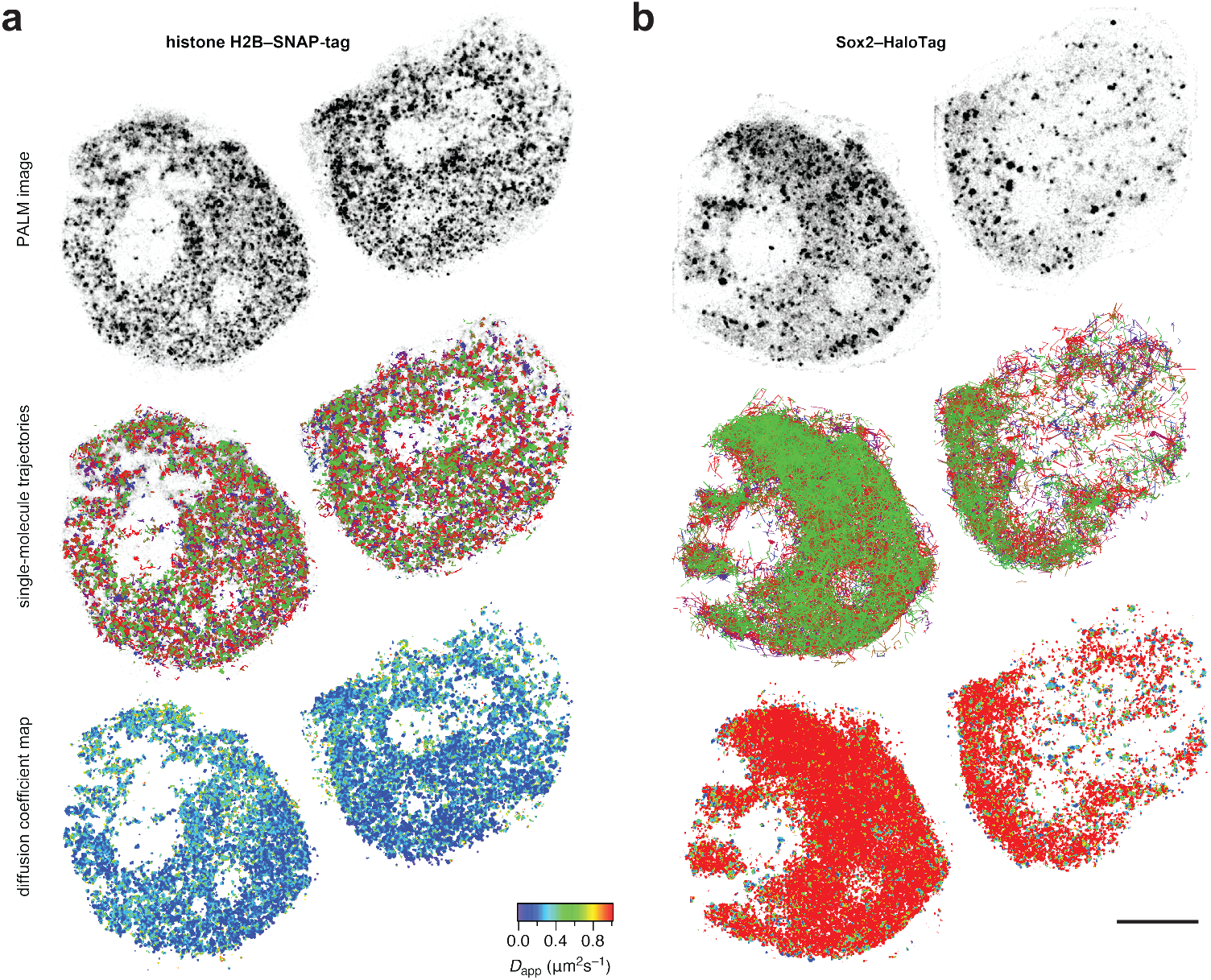
Additional data for simultaneous two-color SPT-PALM experiment. Two live ES cells expressing histone H2B–SNAP-tag labeled with PA-JF_646_-SNAP-tag ligand (**9**) and Sox2–HaloTag labeled with PA-JF_549_-HaloTag ligand (**5**). Upper images show localization microscopy image (PALM), center images show cumulative single-particle trajectories, and lower images show apparent diffusion coefficient map. Scale bar: 5 μm. (**a**) Images from histone H2B–SNAP-tag labeled with **9**. (**b**) Images from Sox2-HaloTag labeled with **5**.

## Supplementary Video Legends

**Supplementary Video 1.** One-photon activation (405 nm) of an ES cell expressing Sox2–HaloTag labeled with PA-JF_549_-HaloTag ligand.

**Supplementary Video 2.** Two-photon activation (800 nm; spelling “HHMI”) in a HeLa cell expressing histone H2B–HaloTag labeled with PA-JF_549_-HaloTag ligand followed by full-field one-photon activation (405 nm).

**Supplementary Video 3.** Side-by-side comparison of 3D single-particle tracking (SPT) experiments in an ES cell expressing Sox2–HaloTag labeled with TMR HaloTag ligand (left) in SPT-*d*STORM mode or PA-JF_549_ (right) in SPT-PALM mode.

**Supplementary Video 4.** One-photon activation (405 nm) of an ES cell expressing GFP–HP1 (green) and Sox2–HaloTag labeled with PA-JF_646_-HaloTag ligand (magenta).

